# The Invisible Heterogeneity of a Forest – Beta Diversity of Volatiles

**DOI:** 10.1101/2025.09.03.673352

**Authors:** Lena Carlson, Jörg Müller, Mirjana Bevanda, Peter Biedermann, Pia M. Bradler, Sebastian Dittrich, Bronwyn Lira Dyson, Andreas Fichtner, Antonio Jose Castañeda-Gomez, Goddert von Oheimb, Kerstin Pierick, Luisa Pflumm, Oliver Mitesser, Julia Rothacher, Clara Wild, Thomas Schmitt

## Abstract

Forest structural heterogeneity affects biodiversity, yet how changes in forest structure influence the spatial patterns of forest chemical heterogeneity remains poorly understood. Volatile organic compounds (VOCs) create invisible chemical landscapes that influence forest ecosystem processes, but whether VOC β-diversity patterns respond to silviculture or disturbance caused heterogeneity remains unknown. We quantified how enhanced structural beta complexity (ESBC) treatments affect VOC β-diversity patterns and investigated potential drivers and ecological effects in temperate production forests. Using the experimental BETA-FOR framework, we sampled ambient forest air at the forest floor and 1 m heights across 234 forest patches in six German regions using Tenax/Carboxen adsorbent traps analyzed via TD-GCMS. Results from generalized linear beta regression models showed that β-diversity of VOCs increased significantly at 1 m height in ESBC forests compared to control forests, but this increase was not significant at the forest floor. In contrast to studies on plants, fungi and animals, the main driver for increasing beta-diversity in VOCs was not the heterogeneity of canopy openness, but the amount of deadwood. Using saproxylic beetles as a test group, we found that saproxylic beetle community dissimilarity increased with VOC dissimilarity, but only for forest floor VOCs. Our finding adds a new component to the framework of habitat heterogeneity, the invisible gradient of volatile diversity utilized by many forest organisms. Furthermore, we provide the first evidence that enhancing the heterogeneity of forests, and particularly of the dead wood, increase not only the structural heterogeneity but also the volatile β-diversity.

## Introduction

Increasing bird diversity with the increasing vertical heterogeneity of forests formed the basis for the habitat heterogeneity hypothesis in the early 1960s (MacArthur & MacArthur, 1961). Since then, several heterogeneity gradients affecting species diversity have been identified, including structural heterogeneity (e.g., horizontal and vertical forest structure, dead wood distribution), topographic heterogeneity, and biotic heterogeneity (e.g., plant diversity) (Heidrich et al., 2020; Stein & Kreft, 2015). However, one heterogeneity gradient that remains largely unexplored due its invisible character is the dissimilarity of the composition of volatile organic compounds. These VOCs are metabolites released by all living organisms and organic matter as byproducts of metabolic processes, creating an invisible chemical scentscape (the volatilome) that influences numerous forest ecosystem processes. The diversity of VOCs may reflect organism diversity: more diverse biological communities produce more heterogeneous volatilomes (Abis et al., 2020; Courtois et al., 2009; Kessler & Kalske, 2018a), while recent studies show the relationship is complex and context dependent due to environmental influences, multitrophic interactions, and scale dependencies (Berkum et al., 2025; Dixon & Dickinson, 2024; Kessler & Kalske, 2018b). Many VOCs have evolved into chemical communication networks that arguably predate auditory and visual systems (Steiger et al., 2011). In forests, VOCs mediate inter- and intra-organism communication, serve as carbon sources for microbes (Ramirez et al., 2010), and influence atmospheric chemistry (Dicke & Baldwin, 2010; Peñuelas & Llusià, J, 2003). For instance, bark beetles use anti-aggregation pheromones to identify when host trees are sufficiently colonized (Frühbrodt et al., 2023), bees use floral volatiles to locate pollen sources (Raguso, 2008), while saproxylic beetles are attracted to chemical signatures of decomposing dead wood (Graf et al., 2021). VOCs establish intricate chemical information networks that function with structural heterogeneity as an important component of forest ecosystems. Understanding this dimension of forest ecosystems is critical since structural changes through disturbance and management may create VOC heterogeneity alongside physical changes, potentially creating invisible heterogeneity gradients.

While vertical VOC heterogeneity is well documented in forest canopies (Petersen et al., 2023; Ringsdorf et al., 2024; Schuman, 2023; Sulzer et al., 2025; Yáñez-Serrano et al., 2018), the horizontal volatile heterogeneity of forest landscapes has rarely been studied.

Current management practices used to enhance structural complexity in otherwise homogeneous production forests include dead wood retention (Großmann et al., 2023; Gustafsson et al., 2012) and canopy gap creation (C. Kern et al., 2016; C. C. Kern et al., 2014; Tong et al., 2024), both of which could potentially influence VOC β-diversity patterns across multiple spatial scales through various drivers. Dead wood may lead to an increase in VOC ß-diversity through decomposition processes that create diverse chemical signatures from different decomposer communities (Mäki et al., 2021), while different dead wood components (standing vs. lying) provide distinct microhabitats (Krumm et al., 2013; Lachat et al., 2019) that may in turn affect VOC emissions. Canopy gaps alter microclimate and understory vegetation composition (Abd Latif & Blackburn, 2010; C. C. Kern et al., 2013), potentially driving VOC β-diversity through effects on plant emissions and decomposition processes. As sample organisms, saproxylic beetle communities may respond to VOC patterns by colonizing chemically distinct substrates and contribute to the volatilome as they themselves release VOCs (Graf et al., 2021; Leather et al., 2014).

Here, we conducted the first landscape-scale field study sampling ambient forest air at the forest floor and 1 m height across 234 patches. We compared 11 experimentally heterogenized forests by increasing the between forest patch heterogeneity (ESBC) via deadwood and gap enhancement to 11 homogenous (Control) forests. Based on these potential influences, we tested the following hypotheses:

H1: Enhanced districts show higher within-district VOC β-diversity than control districts, due to an increase in (H1a) heterogeneity of deadwood volumes, and (H1b) heterogeneity of canopy cover. H2: Dead wood type affects VOC β-diversity, with lying dead wood having stronger effects than standing dead wood, and patches with both types having the strongest effects. H3 Saproxylic beetle communities respond to VOC β-diversity patterns.

## Methods

### Experimental Design and Site Description

This study was conducted on 234 structurally-manipulated forest patches within the BETA-FOR experimental framework across six regions (University Forest, Bavarian Forest, Passau, Hunsrück, Saarland, Lübeck) spanning approximately 900 km^2^ in Germany (Müller et al., 2022). The experimental design compares 11 district pairs of enhanced structural beta complexity (ESBC) districts and control districts. Four different treatments were implemented on forest patches in the enhanced districts including experimental dead wood addition (logs and snags, logs, snags, and stumps) and canopy gap creation. Treatments were implemented either spatially aggregated or distributed resulting in a total of eight treatment patches and one control patch per enhanced district with nine corresponding control patches in the control district (**Figure 1**). An exception was made on University Forest patches, where additional treatments led to 14 different configurations. Preliminary analyses showed no effect of spatial aggregation on VOC patterns, so this factor was excluded from all subsequent analyses.

**Figure 1:**
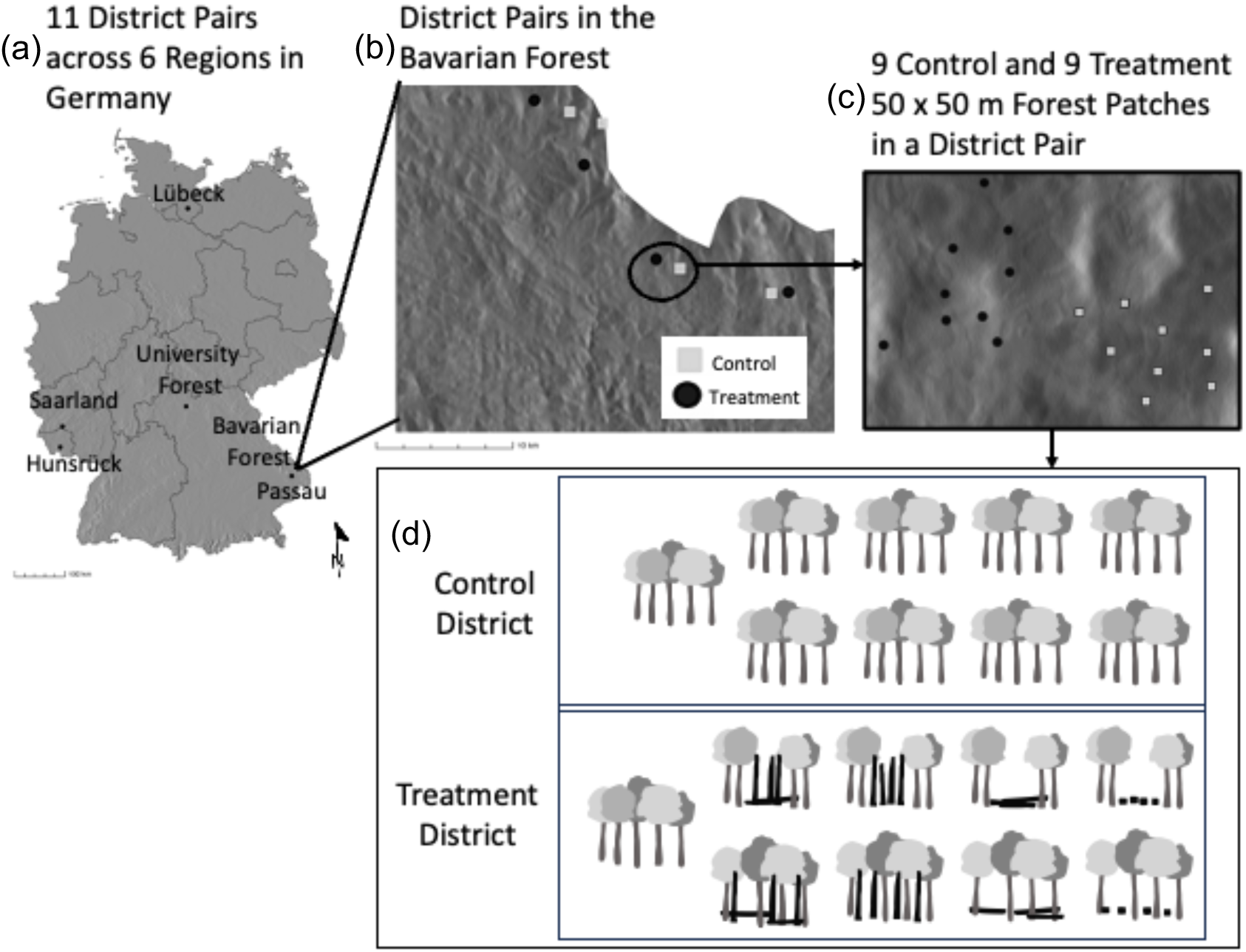
The nested BETA-FOR experimental design spans 234 forest patches in 11 district pairs in six regions across Germany: Passau, Bavarian Forest, University Forest, Hunsrück, Saarland and Lübeck (a and b). Each district pair contains structural beta complexity (ESBC) treatments and controls (c). Treatment districts contain 9 patches (50 × 50 m). Control districts have a homogeneous canopy and minimal dead wood (d, top row) versus enhanced districts with varied canopy structure, gaps, and diverse dead wood components (d, bottom row).

We sampled 234 forest patches (50 × 50 m) in 11 district pairs of control and treatment patches to assess how structural heterogeneity affects volatile organic compounds in German beech production forests. After excluding faulty samples, we analyzed 230 samples from 1 m height and 232 from forest floor level.

### VOC Sample Collection

Volatile organic compounds (VOCs) were collected from 234 BETA-FOR forest patches throughout July 2023. Sampling was conducted simultaneously at two heights at the center of each forest patch: 1 m above ground level and at ground level—forest floor—to capture possible vertical variation in VOC emissions (**Figure 2, a**). VOC samples were collected using conditioned quartz glass tubes (15 mm × 1.9 mm inner diameter) packed with Tenax and Carboxen adsorbents (1.5 mg each) and secured with glass wool plugs (**Figure 2, b**). The sampling tubes were connected to battery-powered DC pumps (Fürgut, Tannheim, Germany), drawing ambient forest air through the tubes at a flow rate of 1.1 L/min for 30 minutes per sample. This sampling protocol was adapted from methods described by Otieno et al. (2023). Due to logistical constraints and given the large spatial scale of this study, we prioritized extensive spatial coverage (234 patches across six regions) over within-patch replication, collecting one sample per height per patch.

**Figure 2:**
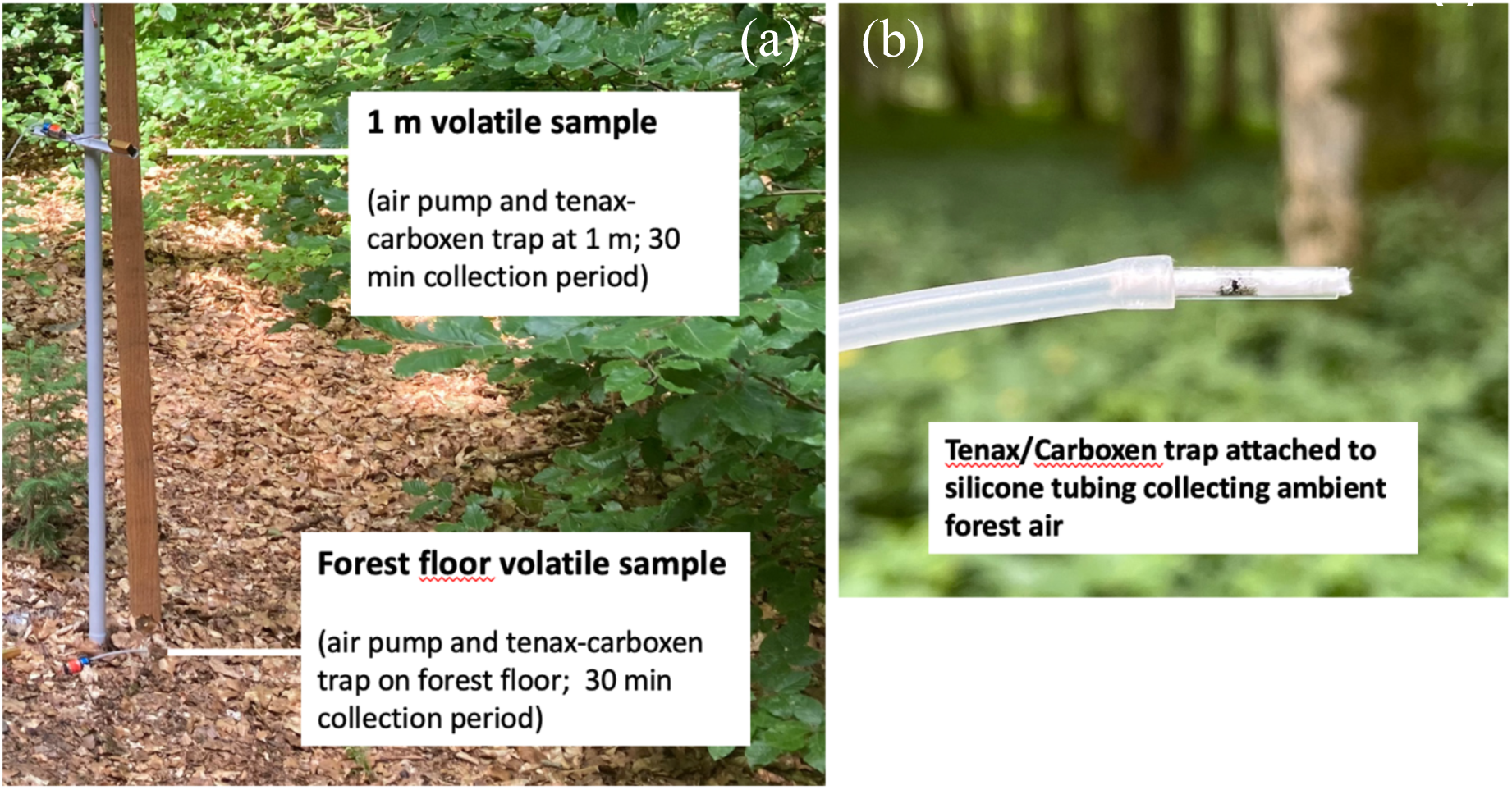
Volatile organic compounds (VOCs) were sampled at 1 m height and the forest floor using Tenax/Carboxen adsorbent traps connected to battery-powered air pumps via silicone tubing. Ambient forest air was sampled for 30 minutes at 1.1 L/min flow rate at the center of each of the 234 BETA-FOR forest patches (Carlson 2025).

Ambient air sampling captures the integrated volatile profile of forest patches—a composite of emissions from multiple sources (e.g., vegetation, decomposing organic matter, soil, soil microbes, fungi, animals) that have mixed in the forest air. This approach allows us to characterize the overall ‘VOC fingerprint’ or volatilome of each forest patch.

To focus the scope of the study and further minimize variation in the samples, we collected samples between the hours of 9:30 and 17:30 during daylight hours when VOC production is highest and on dry days without rain/precipitation—since rain leads to a short-term increase in VOC emissions (Greenberg et al., 2012; Lappalainen et al., 2009) and to ensure the functionality of our VOC pumps—and throughout July.

### Tenax/Carboxen VOC Trap Adsorbents

The VOC traps contained an equal mix (1.5 mg each) of Tenax and Carboxen adsorbents to cover a wider range of VOCs in the forest ecosystem. Tenax leads to sample losses for smaller, more volatile compounds but is effective for larger compounds (Pollmann et al., 2005), and Carboxen is especially efficient for sampling smaller more volatile compounds (Schieweck, 2018). By combining both, we ensure sampling of a wider range of compounds in a highly diverse ecosystem setting.

### Chemical Analysis and Data Processing

VOC samples were analyzed by thermal desorption gas chromatography-mass spectrometry (TD-GC/MS) using a Markes TD100-xr thermal desorption (Markes, Offenbach am Main, Germany) unit coupled to an Agilent 7890B/5977 GC-MS system (Agilent Technologies, Palo Alto, USA). For the chemical analysis, each glass tube containing the Tenax/Carboxen sorbent was transferred to sorption tubes, which were then loaded into the thermal desorber. In the thermal desorber, the sorption tubes were heated in a stream of nitrogen carrier gas to release VOCs from the sorbent materials using the following parameters: flow rate 20 ml / min, flow temperature 180°C, minimum delivery pressure 2 psi; pre-desorption with dry purge time 10 min and purge flow 40 ml / min; tube desorption for 10 min at 260°C with trap flow 40 ml/min. The desorbed VOCs were then directed to a Peltier-cooled focusing trap, which was rapidly heated in the counter-current of carrier gas to inject the VOCs into the GC column. Chromatographic separation was achieved using a HP-5MS UI capillary column (30 m x 0.25 ID; df = 0.25 µm, Agilent Technologies, Palo Alto, USA) with helium as the carrier gas (flow rate 1.287 ml/min, at a constant pressure of 1 bar). The temperature program started at 40°C and increased at 5°C/min to 300°C, with a total run time of 56 min per sample. The TD-GC/MS methods were adapted from Otieno et al. (2023).

Raw chromatographic data were integrated using Agilent MassHunter software to quantify individual VOC peaks, with each peak corresponding to one compound (see Figure S *1*: Representative gas chromatography profile showing the VOC complexity of forest air in a single forest floor sample, with individual volatile compounds (each peak corresponds to one compound) spanning from highly volatile (left) to semi-volatile (right) and range from monoterpenes to GLVs, sesquiterpenes, esters, humic-derived compounds to phenolic compounds as major compound classes. in supplements for an example of a chromatogram and corresponding compound classes). Peak alignment across samples was performed using the GCalignR package (Ottensmann et al., 2018) to ensure consistent compound quantification across all samples. This processing workflow generated two data outputs for each detected compound: mean retention time and peak height measurements. Individual compounds were not identified to specific VOC classes, as aligned peak quantification was sufficient for analyzing VOC β-diversity patterns in forest ecosystems.

While background compounds and potential instrument contamination were not filtered from the data, experimental procedures were randomized throughout, and chemical noise was equally present across all samples.

### VOC Data Structure

The complete VOC dataset contained 724 peaks detected at 1 m height and 727 peaks at the forest floor level. Most peaks (43%) appeared in more than half of all samples, indicating consistent detection across patches, while only 7.4% and 7.6% of compounds, respectively, were rare (detected in fewer than 5% of samples). The top 10% most frequently detected compounds accounted for 21% of all detections, showing that no single compounds dominated the chemical profiles. This balanced distribution of common and rare compounds provided the chemical diversity needed for β-diversity analyses across all forest patches (see Figure S 2 in supplements for VOC distribution plots).

### Environmental Variables

Average air temperature was calculated for each sample using EasyLog logger temperature data recorded at patch centers within ±30 minutes of the 30-minute VOC sampling period, since exact time periods were not available for all sampling periods. Tree species composition surveys and basal area measurements were conducted on all patches (Pierick & Ammer, 2025). Dead wood volume and type (including stumps, logs, snags, and habitat trees) were surveyed on all patches (Junginger et al., 2025). 278 herbaceous and woody species up to 1 m height were recorded on five 4 m radius sub patches per patch during the vegetation period in 2023 (Bradler & Fichtner, 2025). Canopy cover was measured by LiDAR via drone flights using the penetration ratio at 6 m above ground (Müller & Brandl, 2009) (M. Bevanda, A.Castaneda-Gomez, L. Pflumm, unpublished data).

Since many fungal sporocarps are closely associated with dead wood we used data on sporocarp presence/absence from stumps, logs and snags surveyed on the BETA-FOR patches in autumn of 2023 and 2024 (Lira Dyson & Bässler, 2025) . Beetles were sampled using two non-attracting flight-interception traps per patch for 2023. Each trap consisted of a crossed pair of transparent plastic shields (40 cm × 60 cm), a plastic roof, and a funnel leading to a bottom plastic container filled with a diluted solution of copper sulfate (3 %) to preserve the caught arthropods. Data was filtered to include only saproxylic beetles for a total of 448 species (Müller, 2025). Data collection followed standardized BETA-FOR protocols established by Müller et al. (2022) to ensure consistency across the 11 pairs.

### Treatment Classification

While the BETA-FOR framework implements nine to 14 structural treatments depending on the region (Control, Stumps, Logs, Snags, Logs and Snags, Total Tree Removed, Crowns, and Habitat Trees) and VOC sampling was conducted across all treatment types, we analyzed using a simplified classification better suited for this large-scale chemical ecology analysis. For within-district heterogeneity analyses, we calculated pairwise comparisons among all patches within each district (treatment patches vs. treatment patches, control patches vs. control patches) to test whether treatment districts were more heterogeneous than control districts. Here we examined continuous gradients of structural complexity based on measured dead wood volume and canopy cover rather than categorical treatment assignments. The continuous variables (dead wood volume, canopy cover) capture the actual structural complexity achieved regardless of intended treatment design and treatments naturally create gradients in these variables (e.g., patches with logs and snags inherently have higher dead wood volumes than stump-only patches). Total dead wood volume included stumps, logs, high stumps, and habitat trees treated as high stumps, but excluded crown material, which had largely decomposed by the time of VOC sampling. This approach allowed us to test both categorical treatment effects (Enhanced vs. Control districts; dead wood type; canopy cover) and continuous structural relationships—based on actual measured complexity (dead wood volume by type, canopy cover).

### Terminology

In this study VOCs are the volatile organic compounds detectable by our TD-GC/MS method, encompassing compounds from C5 to just over C40, compounds containing 5 to 40 carbon atoms. While VOCs are responsible for the scent of the forest, we refrained from using scent-related terminology, preferring the use of “VOCs”, “VOC β-diversity” or “chemical compounds” when referring to the compounds. When referring to the total VOCs present at a forest patch or district, we chose “volatile profile” or “volatilome” over “scentscape” for consistency. The volatilome is the totality of VOCs detected at a defined time and area, in this study referring to the collective VOC profile of forest patches, forest districts, or the broader forest ecosystem.

### Statistical Analysis

All statistical analyses were performed in R version 4.4.3 (R Core Team, 2025). The tidyverse package (Wickham et al., 2019) was used for data handling and visualization. We conducted two analyses to address different aspects of VOC heterogeneity:

**(1) Analysis of within-district heterogeneity:** To test treatment effects, we compared VOC β-diversity between treatment and control forest districts. Within-district heterogeneity was quantified by calculating pairwise VOC dissimilarity (True Jaccard distance) between all possible patch combinations within each district, resulting in a distribution of within-district dissimilarity values for each district. We then tested whether treatment districts showed higher within-district heterogeneity than control districts using beta regression generalized linear mixed models (GLMMs) implemented in glmmTMB (Brooks et al., 2017), with district type (treatment vs. control) as a fixed effect and random effects for region, sampling hour, and date. Pairwise dissimilarity values were adjusted using a small constant (0.00001) to accommodate the beta distribution requirements.
**(2) Analysis of environmental drivers**: To identify drivers underlying VOC patterns, we examined which environmental factors drive chemical dissimilarity between any two patches across the landscape using Multiple Regression on Distance Matrices (MRM) analyses (999 permutations) from the ecodist package (Goslee & Urban, 2007). This analysis included all 234 patches regardless of district assignment to maximize the statistical power to detect environmental relationships. Spatial autocorrelation in VOC patterns was also addressed using the MRM analysis. The significant effect of spatial distance (p < 0.001) confirmed spatial structure in our data. This was expected since our study spanned six regions across 900 km², and patches that are geographically farther apart are likely to differ in environmental conditions (climate, elevation, local species composition) that influence VOC composition.

VOC samples were analyzed separately by sampling height (1 m and forest floor) to capture possible height-specific responses, as differences in forest air composition have been found particularly in the forest canopy and close to the soil (Mäki et al., 2019; Noe et al., 2012). Fewer studies have been conducted on ambient understory air (Isidorov & Jdanova, 2002; Jüttner, 1986).

## Results

Our findings show ESBC treatments significantly increased within-district VOC β-diversity at 1 m height (effect size = 0.151, p < 0.001) but not at the forest floor (effect size = 0.019, p = 0.122) (**Figure 3**), demonstrating that structural enhancement creates VOC heterogeneity within temperate, beech-dominated production forests (**Error! Reference source not found.**).

**Figure 3:**
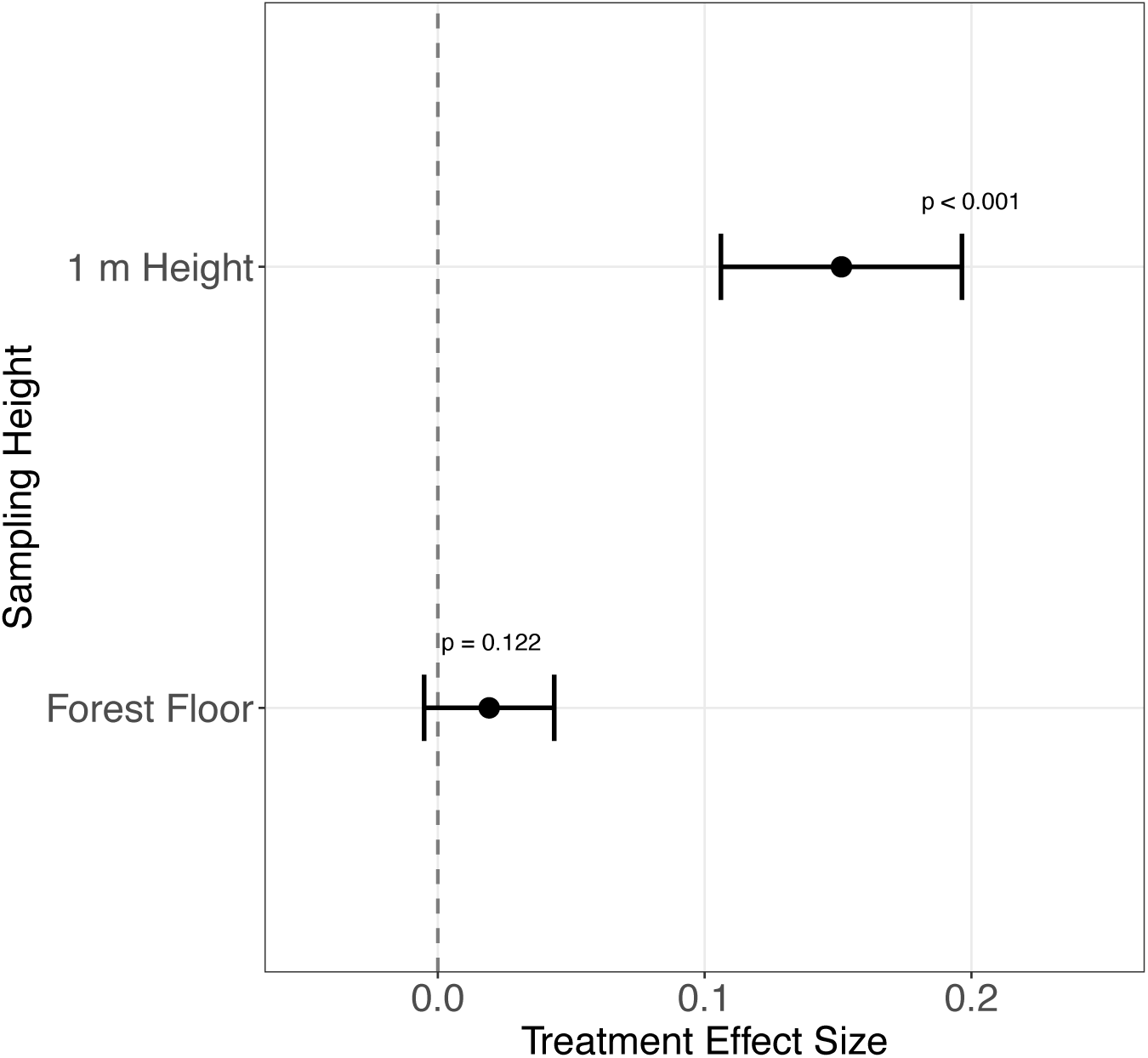
Enhanced structural beta complexity (ESBC) treatments increase within-district VOC β-diversity at 1 m height but not at the forest floor. Treatment districts show significantly higher within-district VOC β-diversity than control districts at 1 m height (p < 0.001) but not at forest floor level (p = 0.122). Effect sizes (β ± 95% CI) from beta regression models demonstrate that structural enhancement creates chemical heterogeneity primarily in ambient forest air rather than through forest floor processes.

Treatment districts showed considerably higher internal chemical heterogeneity than control districts, but only at 1 m height. These height-specific responses show that treatment districts develop internal VOC heterogeneity that distinguishes them from more homogeneous control districts.

To understand the drivers underlying these district-level patterns, we examined environmental and treatment-related drivers of VOC dissimilarity between individual patches across the entire landscape (**Figure 4**). While enhanced districts showed greater internal heterogeneity at 1 m height, a closer look at the potential drivers revealed that forest floor processes actually explained more variance in VOC patterns when comparing patches across all districts and regions.

**Figure 4:**
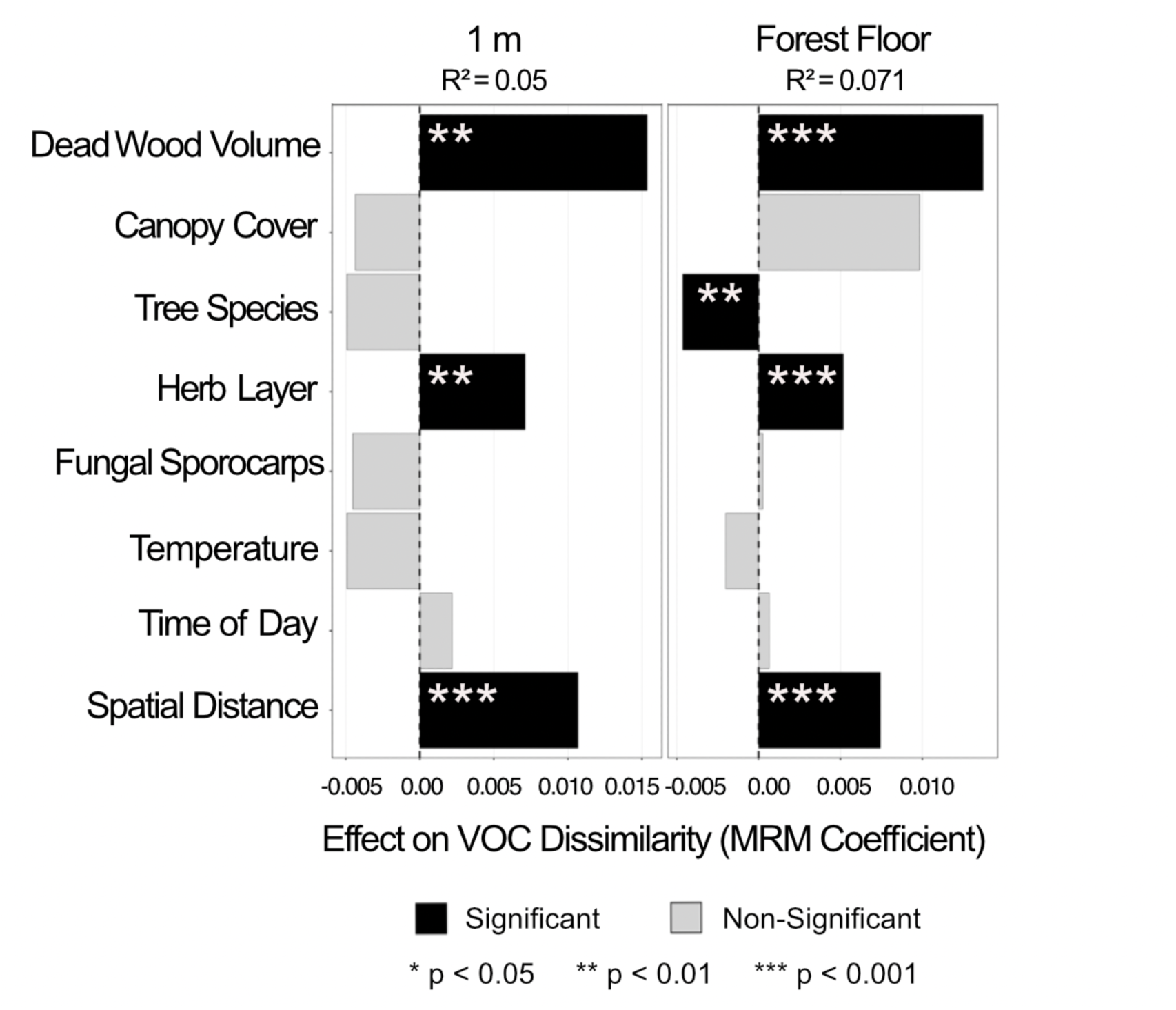
Environmental drivers of VOC dissimilarity between forest patches from MRM analysis. Significant effects (black bars, p < 0.05) show dead wood volume as the primary treatment-related driver, with additional effects from herb layer, spatial distance, and tree species diversity (significant negative effect at forest floor). Effect sizes indicate the magnitude and direction of influence on pairwise VOC dissimilarity. Longer bars = stronger effects on chemical dissimilarity.

Dead wood volume emerged as the primary treatment-related driver of VOC heterogeneity at both sampling heights (1 m: effect size = 0.015, p < 0.005; forest floor: effect size = 0.014, p < 0.001), while canopy cover showed no significant effects despite creating microclimatic gradients visible on the patches (**Figure 4**). The herb layer significantly influenced VOC patterns at both heights (1 m: p < 0.004; forest floor: p < 0.001). Tree species diversity showed an unexpected pattern: no effect at 1 m height but a significant negative effect on VOC dissimilarity at the forest floor (p < 0.003). Spatial distance showed the strongest overall effect across both heights (p < 0.001). Temperature and time of day were not significant. Fungal sporocarps also showed no significant effects on VOC patterns. The MRM explained more variance at the forest floor (R² = 0.071) than at 1 m height (R² = 0.05).

When considering dead wood diversity and dead wood volume, volume alone had no significant effect on VOC dissimilarity at either sampling height (**Figure *5***). However, patches containing both standing and lying dead wood objects significantly increased VOC heterogeneity at both 1 m height (p < 0.002) and forest floor (p < 0.001), showing that component diversity creates chemical complexity beyond simple volume effects.

**Figure 5:**
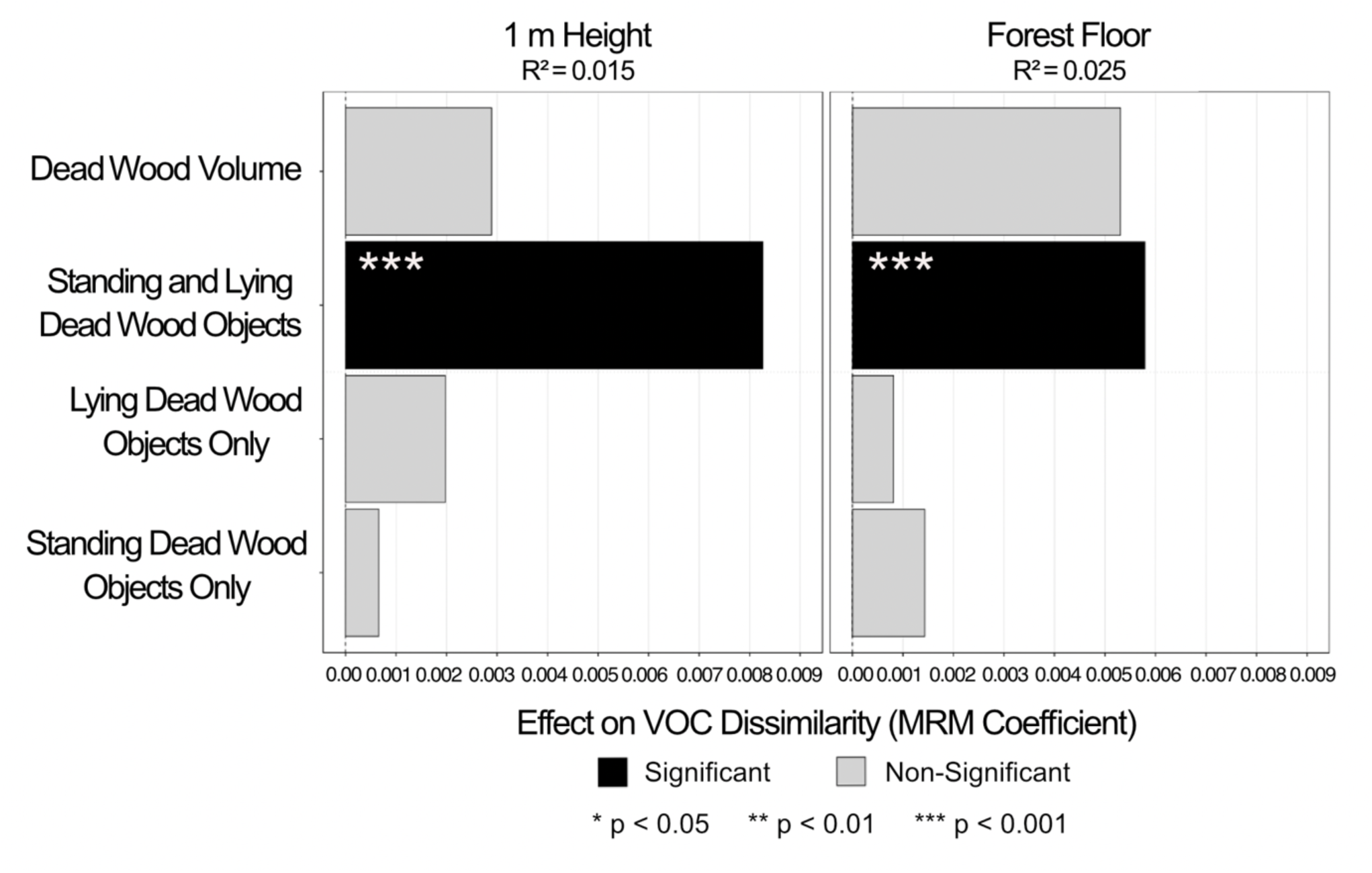
Dead wood diversity enhances VOC heterogeneity beyond volume alone. A comparison of continuous (dead wood volume) versus categorical (object types) predictors of VOC dissimilarity shows only patches with both standing and lying dead wood components significantly increase VOC heterogeneity, demonstrating that component diversity creates chemical complexity beyond simple volume effects. Bar length shows effect size, with significant effects in black (p < 0.05). Longer bars = stronger effects on chemical dissimilarity.

Individual dead wood object types had no significant effects meaning standing dead wood only and lying dead wood only patches did not significantly differ from controls in their VOC heterogeneity patterns. The MRMs explained more variance at the forest floor (R² = 0.025) compared to 1 m height (R² = 0.015).

Forest floor VOC β-diversity significantly influenced saproxylic beetle species richness (q0, p<0.02) but had no significant effects on Shannon (q1) or Simpson (q2) diversity measures (**Figure 6**) In contrast, 1 m height VOCs showed no significant effects on saproxylic beetle communities across any diversity order.

**Figure 6:**
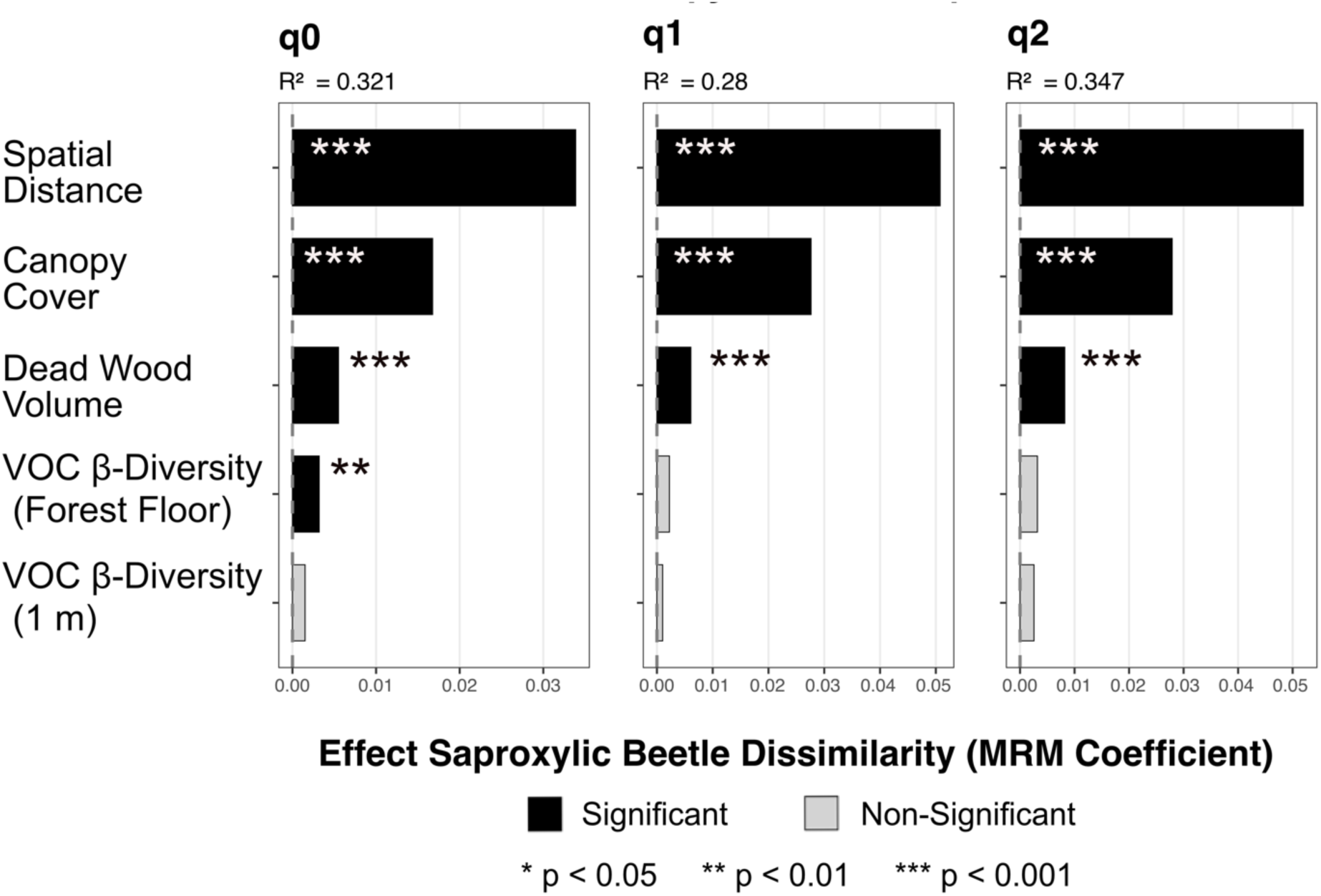
Forest floor VOC β-diversity influences saproxylic beetle species richness. Effects of VOC β-diversity and structural variables on beetle community dissimilarity across diversity orders (q0-q2). Forest floor VOC patterns significantly influence beetle species richness (q0) but not Shannon (q1) or Simpson (q2) diversity measures, while 1 m height VOC patterns show no significant effects across any diversity order. Spatial distance, canopy cover, and dead wood volume consistently drive beetle community structure across all diversity measures. Bar length shows effect magnitude, with significant effects in black (p < 0.05). Longer bars = stronger effects on chemical dissimilarity.

Structural variables consistently drove beetle community patterns across all diversity measures. Spatial distance showed the strongest effects across diversity orders (q0-q2, all p < 0.001), followed by canopy cover and dead wood volume (all p < 0.001).

## Discussion

While vertical VOC heterogeneity in forests has been well documented in canopy studies (Petersen et al., 2023; Ringsdorf et al., 2024; Sulzer et al., 2025; Yáñez-Serrano et al., 2018), the horizontal heterogeneity of volatiles across forest landscapes has received little attention (Hagiwara et al., 2024). This study represents the first landscape-scale investigation of VOC β-diversity patterns, comparing chemical heterogeneity between 11 ESBC and 11 control districts to determine whether enhanced management can create spatial chemical gradients alongside structural changes.

This patch-level landscape-scale study required several methodological considerations: Our sampling across 234 forest patches generated highly skewed VOC data with frequent zeros and peak height values ranging from zero to 296450937.22 counts, representing the signal intensity per time point detected by the mass spectrometer for each compound. We therefore analyzed presence/absence data rather than abundance, which proved effective for detecting β-diversity while avoiding the statistical issues associated with extreme data distributions.

Despite 30-minute sampling representing only snapshots of volatile diversity—as VOC emissions can vary greatly with environmental conditions, time of day, and organisms present—treatment effects were detectable across this spatial scale, demonstrating that ESBC treatments create measurable changes in forest volatilomes (**Figure 3**).

Spatial distance significantly influenced VOC dissimilarity patterns, which was expected given that patches separated by greater distances are exposed to different environmental conditions across the 900 km² study area.

Neither sampling time nor temperature had significant effects on our results, which proved advantageous since we had to sample throughout the day to cover 234 patches across Germany. While VOCs typically follow diurnal emission patterns with higher daytime emissions and respond to temperature, light, humidity, and rainfall (Greenberg et al., 2012; Monard et al., 2021; Schuman et al., 2016), our treatment effects were unaffected by these potential confounding factors.

Our results suggest that ESBC treatments create chemical heterogeneity within forest landscapes, generating a patchwork of differing volatilomes (**Figure 3**). The height-specific response may reflect differences in VOC mixing through environmental factors such as wind, canopy influence, proximity to understory vegetation, decomposition processes such as dead wood and litter and soil emissions. At the forest floor, more homogeneous emissions from litter decomposition and soil processes, combined with stable microclimatic conditions, may create more uniform VOC responses across patches despite ESBC treatments.

Our results indicate that management can create heterogeneous volatilomes across multiple spatial scales, creating invisible heterogeneity gradients with potential ecological consequences for species communities such as saproxylic beetles navigating forest landscapes (Randlkofer et al., 2010).

Further analysis of drivers across this landscape-scale study reveals multiple drivers creating VOC heterogeneity (**Figure 4**): dead wood volume emerges as the primary treatment-related driver at both heights, while canopy cover shows no significant effects despite creating microclimatic gradients and most likely influencing herb layer growth. This suggests that decomposition processes may emit more prominent VOC signals in temperate beech forest understories than light-driven emission responses from increased vegetation due to changes in canopy cover, supporting the height-stratification we see in our district-level results (**Error! Reference source not found.**).

Herb layer coverage significantly influenced VOC patterns, indicating that understory vegetation contributes to landscape-scale chemical heterogeneity despite limited research on understory plant emissions (Isidorov et al., 2022). Tree species diversity showed unexpected negative effects at forest floor level, contrasting with canopy-level diversity patterns where different species emit distinct VOC profiles (Antonelli et al., 2020), possibly reflecting soil-mediated homogenization processes.

In contrast to recent studies showing that canopy openness predominantly drives animal, plant and fungi responses to ESBC treatments on BETA-FOR patches (Massó Estaje et al., 2025; Rothacher et al., 2023), dead wood emerged as the main driver of VOC β-diversity (**Figure 4**). While canopy gaps can impact light availability, temperature and solar radiation (Muscolo et al., 2014), and VOC emissions can be affected by these factors (Maja et al., 2016; Owen et al., 2002), dead wood volume was the sole significant treatment-related driver of chemical heterogeneity. This suggests that decomposition and microbial processes, rather than light-driven changes, primarily drive the observed VOC β-diversity patterns at least at the strata up to 1 m height form the forest floor. This aligns with recent findings that microbial activity and decomposition are key regulators of forest VOC dynamics (Lee et al., 2025). Our findings that VOCs are more affected by deadwood than by canopy density further demonstrates that VOCs form a rather independent heterogeneity gradient to the horizontal heterogeneity in forests dominated by light conditions (Heidrich et al., 2020). This independence makes this gradient promising to disentangle the heterogeneity in the volatilome versus the physiognomic heterogeneity of forests.

While we predicted lying dead wood in the beech-dominated forests would have stronger effects than standing dead wood on VOC heterogeneity due to increased soil contact and moisture retention, higher beetle diversity and more fungal sporocarps (Uhl et al., 2022), we acknowledge that equally compelling arguments exist for standing dead wood. Our prediction was partly based on our chosen sampling design, with collection heights closer to the ground where processes associated with lying dead wood may have a greater influence. Both deadwood types present distinct advantages that could drive VOC heterogeneity. Neither laying nor standing dead wood objects alone significantly increased VOC β-diversity.

Dead wood diversity rather than individual types drives VOC heterogeneity, suggesting that different microhabitat types generate greater heterogeneity (**Figure *5***). This diversity effect remained significant even when controlling for total dead wood volume, indicating that patches containing both standing and lying dead wood create enhanced VOC heterogeneity through synergistic rather than additive effects. The finding that standing or lying dead wood alone did not have significant effects may reflect very small effect sizes, which statistical methods commonly used in ecology often fail to detect when working with VOCs (Kempraj 2025).

The combined effect of both dead wood types may lead to more heterogeneous volatilomes because of increased organism diversity on forest patches. Although there is an overlap in organisms inhabiting these dead wood types, some species respond to very specific niches and are indicators of either standing or lying dead wood (Graf et al., 2021). Standing dead wood often provides drier, more sun-exposed conditions, while lying dead wood maintains higher moisture content through direct contact with the forest floor. These contrasting environments attract different microbes, fungi and saproxylic beetles (Löfroth et al., 2023). Since deadwood VOC emissions depend on abiotic conditions, chemical processes, and decomposer organisms (Holighaus & Schütz, 2006), standing versus lying deadwood would be expected to create different volatilome signatures since they fill different niches in forest ecosystems (Bauhus et al., 2018). However, such differences may require compound-specific analyses to detect, since our presence/absence approach may lack the resolution to capture subtle compositional variations.

We used saproxylic beetles as a test community to relate to our VOC pattern, as they are known to respond to as well as emit VOCs in host finding, within species communication and habitat identification. Here forest floor VOC β-diversity significantly influenced saproxylic beetle species richness, showing that the patterns we detected have ecological effects (**Figure 6**). This finding is consistent with the reliance of saproxylic beetles on chemical cues to locate deadwood and associated fungi (Holighaus et al., 2014; Holighaus & Schütz, 2006; Leather et al., 2014). This suggests that beetles respond to chemical differences between patches at the forest floor level, where treatment districts did not create detectable VOC heterogeneity.

## Broader Implications

This study shows that forest management creates chemical heterogeneity alongside structural changes, expanding habitat heterogeneity theory to include invisible chemical gradients. VOCs represent an ’invisible’ heterogeneity gradient that operates alongside established structural and biotic gradients across landscape scales. Our findings complement documented BETA-FOR treatment effects (Asch, 2025; Kacic et al., 2025; Massó Estaje et al., 2025; Rothacher et al., 2023; Schwarz, 2025) and ecosystem multifunctionality research (Chao et al., 2024), demonstrating that structural enhancement creates effects across multiple scales and ecological components. This study shows forest management can manipulate VOC β-diversity patterns, though the underlying mechanisms need further study.

## Limitations

Our detected treatment effects on VOC beta diversity may represent conservative estimates of the true volatilome differences between enhanced and control districts since we only sampled for an arbitrary 30 minutes at each height per patch. These results clearly demonstrate that dead wood volume and diversity drives VOC compositional changes and increased β-diversity, but we can only speculate about the underlying mechanisms. The increased heterogeneity could result from dead wood adding new decomposition-derived compounds (Holighaus & Schütz, 2006), or creating spatially heterogeneous microhabitats with distinct volatilomes. While our study focused on volatilome-level patterns rather than specific compounds, this approach captures the complex chemical environment that organisms actually experience. Future compound and source-specific analyses will be necessary to truly understand the mechanisms underlying these landscape-scale patterns.

## Conclusion

This study is the first landscape-scale analysis of forest VOC β-diversity responses to structural enhancement, showing that forest management creates an invisible horizontal VOC heterogeneity gradient alongside the physical structural heterogeneity. ESBC treatments significantly increased within-district VOC β-diversity at 1 m height but not at the forest floor, indicating height-specific processes drive these heterogeneity patterns and aligning with studies on the vertical stratification of VOCs. Dead wood diversity, rather than volume alone, emerged as the primary driver of VOC heterogeneity, while saproxylic beetles responded significantly to forest floor VOC patterns. These findings indicate that structural enhancement generates chemical heterogeneity with ecological effects, extending the habitat heterogeneity theory to include invisible dimension of the forest volatilome. Future research should focus on identifying the specific mechanisms driving these responses and examine how individual species contribute to the observed pattern VOC sources, compound classes and–where possible–compounds. Understanding how enhancing forest structure creates these VOC gradients can be used to inform forest management strategies. These findings point to the potential of structural treatments and disturbance to shape β-diversity in forests by influencing both the visible physical and invisible chemical dimensions of habitat heterogeneity.

## Data Availability

The datasets generated and analyzed in this study are available in the Zenodo repository: 10.5281/zenodo.14975137. R code for statistical analyses is available upon request from the corresponding author.

## Acknowledgements

We are grateful to the Deutsche Forschungsgemeinschaft (DFG) for funding the BETA-FOR (FOR5375, 459717468) research unit. Thank you to Lena Rabenhofer for her help in the field and Lena Unterbauer for processing the samples.

---

**Figure S1:**
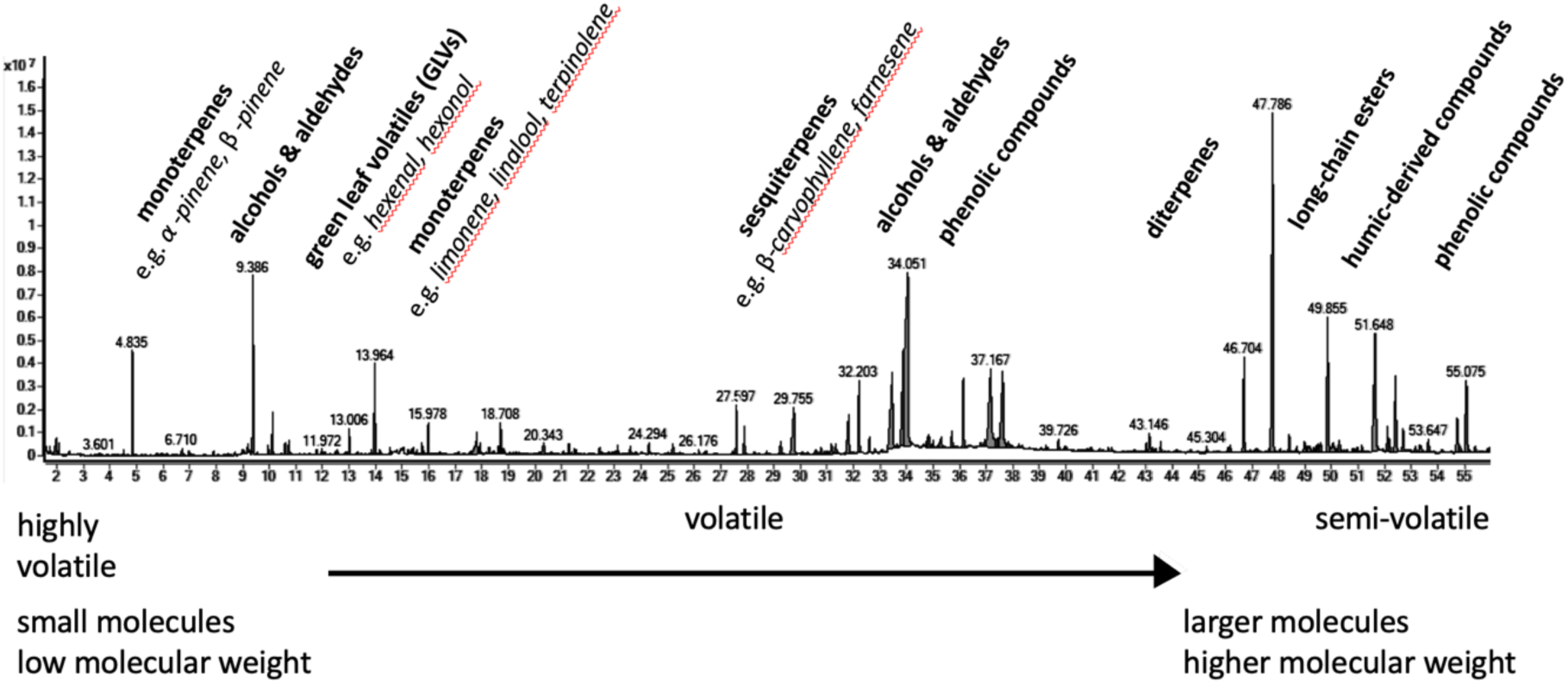
Representative gas chromatography profile showing the VOC complexity of forest air in a single forest floor sample, with individual volatile compounds (each peak corresponds to one compound) spanning from highly volatile (left) to semi-volatile (right) and range from monoterpenes to GLVs, sesquiterpenes, esters, humic-derived compounds to phenolic compounds as major compound classes.

**Figure S2:**
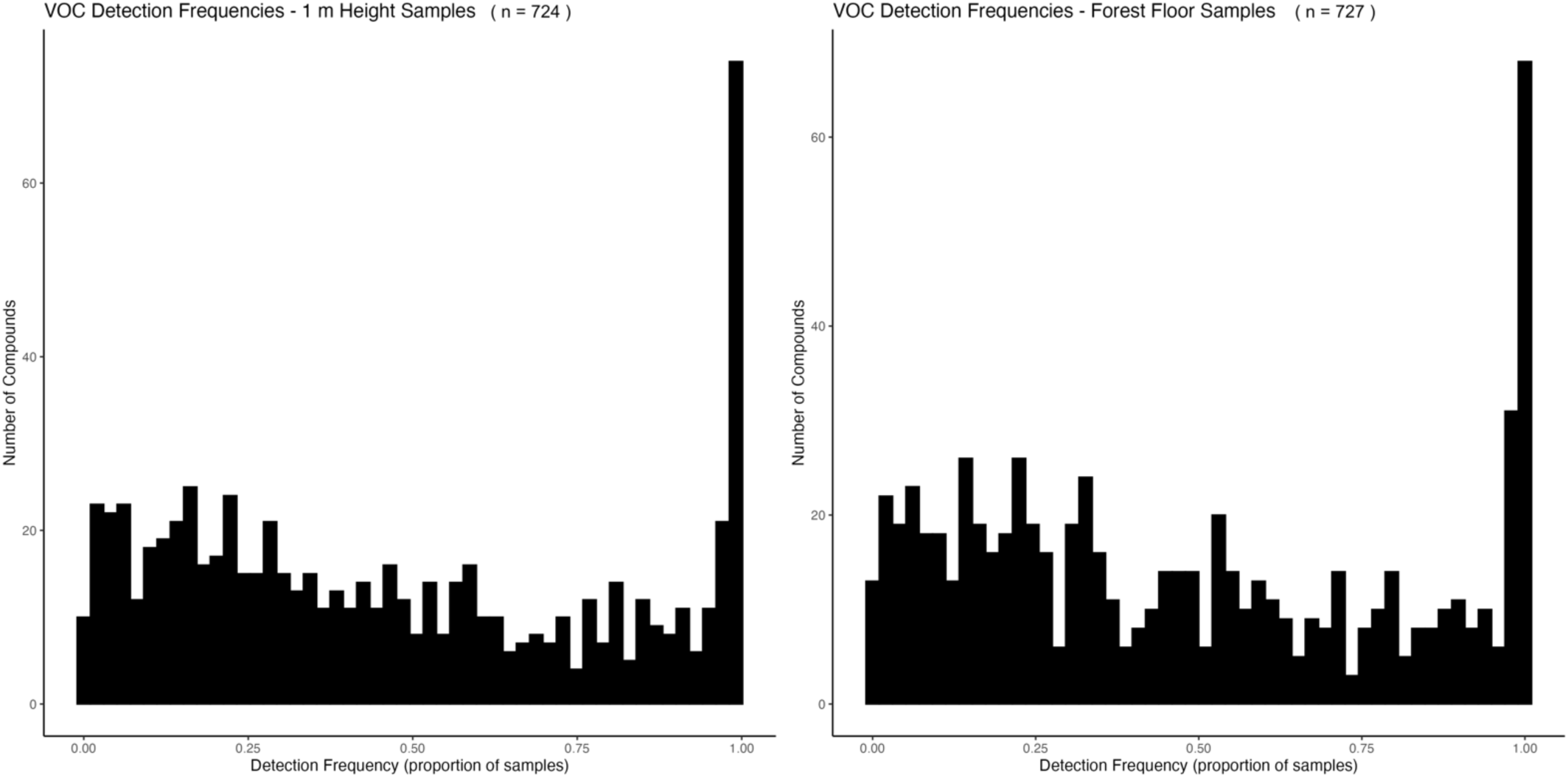
Distribution of VOC frequencies by sampling height. Histograms show the proportion of samples in which each compound was detected for 1 m height samples (n = 724 compounds) and forest floor samples (n = 727 compounds). Most compounds had intermediate detection frequencies, with relatively few compounds detected in either very few (< 10%) or close to all (> 95%) samples. The prominent peak at 1.0 indicates that many compounds were detected consistently across nearly all samples, while other compounds show more variable detection across samples.

## References

Abd Latif, Z., & Blackburn, G. A. (2010). The effects of gap size on some microclimate variables during late summer and autumn in a temperate broadleaved deciduous forest. International Journal of Biometeorology, 54(2), 119–129. 10.1007/s00484-009-0260-1

Abis, L., Loubet, B., Ciuraru, R., Lafouge, F., Houot, S., Nowak, V., Tripied, J., Dequiedt, S., Maron, P. A., & Sadet-Bourgeteau, S. (2020). Reduced microbial diversity induces larger volatile organic compound emissions from soils. Scientific Reports, 10(1), 6104. 10.1038/s41598-020-63091-8

Antonelli, M., Donelli, D., Barbieri, G., Valussi, M., Maggini, V., & Firenzuoli, F. (2020). Forest Volatile Organic Compounds and Their Effects on Human Health: A State-of-the-Art Review. International Journal of Environmental Research and Public Health, 17(18), 6506. 10.3390/ijerph17186506

Asch, J. (2025). *Dung beetles do not profit from enhanced spatial heterogeneity in production forests: A large-scale forest manipulation experiment*.

Berkum, P. M., Albracht, C., Bröcher, M., Solbach, M. D., Stein, G., Bonkowski, M., Buscot, F., Heintz-Buschart, A., Ebeling, A., Eisenhauer, N., El-Madany, T. S., Huang, Y., Kuebler, K., Meyer, S. T., Gershenzon, J., & Unsicker, S. B. (2025). Plant diversity shapes plant volatile emission differently at the species and community level (p. 2025.04.30.651392). bioRxiv. 10.1101/2025.04.30.651392

Bradler, P. M., & Fichtner, A. (2025). *BETA-FOR_SP8_understory_vegetation_surveys_2016-24 (1.0)* [Dataset]. Zenodo. 10.5281/zenodo.14982725

Brooks, M., E., Kristensen, K., Benthem, K., J. ,van, Magnusson, A., Berg, C., W., Nielsen, A., Skaug, H., J., Mächler, M., & Bolker, B., M. (2017). glmmTMB Balances Speed and Flexibility Among Packages for Zero-inflated Generalized Linear Mixed Modeling. The R Journal, 9(2), 378. 10.32614/RJ-2017-066

Courtois, E. A., Paine, C. E. T., Blandinieres, P.-A., Stien, D., Bessiere, J.-M., Houel, E., Baraloto, C., & Chave, J. (2009). Diversity of the volatile organic compounds emitted by 55 species of tropical trees: A survey in French Guiana. Journal of Chemical Ecology, 35(11), 1349–1362. 10.1007/s10886-009-9718-1

Dicke, M., & Baldwin, I. T. (2010). The evolutionary context for herbivore-induced plant volatiles: Beyond the ‘cry for help.’ Trends in Plant Science, 15(3), 167–175. 10.1016/j.tplants.2009.12.002

Dixon, R. A., & Dickinson, A. J. (2024). A century of studying plant secondary metabolism—From “what?” to “where, how, and why?” Plant Physiology, 195(1), 48–66. 10.1093/plphys/kiad596

Frühbrodt, T., Du, B., Delb, H., Burzlaff, T., Kreuzwieser, J., & Biedermann, P. H. W. (2023). Know When You Are Too Many: Density-Dependent Release of Pheromones During Host Colonisation by the European Spruce Bark Beetle, Ips typographus (L.). Journal of Chemical Ecology, 49(11), 652–665. 10.1007/s10886-023-01453-y

Goslee, S. C., & Urban, D. L. (2007). The **ecodist** Package for Dissimilarity-based Analysis of Ecological Data. Journal of Statistical Software, 22(7). 10.18637/jss.v022.i07

Graf, M., Lettenmaier, L., Müller, J., & Hagge, J. (2021). Saproxylic beetles trace deadwood and differentiate between deadwood niches before their arrival on potential hosts. Insect Conservation and Diversity, 15. 10.1111/icad.12534

Greenberg, J. P., Asensio, D., Turnipseed, A., Guenther, A. B., Karl, T., & Gochis, D. (2012). Contribution of leaf and needle litter to whole ecosystem BVOC fluxes. Atmospheric Environment, 59, 302–311. 10.1016/j.atmosenv.2012.04.038

Großmann, J., Carlson, L., Kändler, G., Pyttel, P., Kleinschmit, J. R. G., & Bauhus, J. (2023). Evaluating retention forestry 10 years after its introduction in temperate forests regarding the provision of tree-related microhabitats and dead wood. European Journal of Forest Research, 142(5), 1125–1147. 10.1007/s10342-023-01581-w

Gustafsson, L., Baker, S. C., Bauhus, J., Beese, W. J., Brodie, A., Kouki, J., Lindenmayer, D. B., Lõhmus, A., Pastur, G. M., Messier, C., Neyland, M., Palik, B., Sverdrup-Thygeson, A., Volney, W. J. A., Wayne, A., & Franklin, J. F. (2012). Retention Forestry to Maintain Multifunctional Forests: A World Perspective. BioScience, 62(7), 633–645. 10.1525/bio.2012.62.7.6

Hagiwara, T., Shiojiri, K., Suyama, Y., Matsuo, A., & Ishihara, M. I. (2024). Volatile-mediated plant–plant communication in natural beech forests. Journal of Plant Interactions, 19(1), 2414103. 10.1080/17429145.2024.2414103

Heidrich, L., Bae, S., Levick, S., Seibold, S., Weisser, W., Krzystek, P., Magdon, P., Nauss, T., Schall, P., Serebryanyk, A., Wöllauer, S., Ammer, C., Bässler, C., Doerfler, I., Fischer, M., Gossner, M. M., Heurich, M., Hothorn, T., Jung, K., … Müller, J. (2020). Heterogeneity–diversity relationships differ between and within trophic levels in temperate forests. Nature Ecology & Evolution, 4(9), 1204–1212. 10.1038/s41559-020-1245-z

Holighaus, G., & Schütz, S. (2006). Odours of wood decay as semiochemicals for Trypodendron domesticum L. (Col., Scolytidae). In *Mitteilungen der deutschen Gesellschaft für allgemeine und angewandte Entomologie* (Vol. 15).

Holighaus, G., Weißbecker, B., von Fragstein, M., & Schütz, S. (2014). Ubiquitous eight-carbon volatiles of fungi are infochemicals for a specialist fungivore. Chemoecology, 24(2), 57–66. 10.1007/s00049-014-0151-8

Isidorov, V. A., Pirożnikow, E., Spirina, V. L., Vasyanin, A. N., Kulakova, S. A., Abdulmanova, I. F., & Zaitsev, A. A. (2022). Emission of volatile organic compounds by plants on the floor of boreal and mid-latitude forests. Journal of Atmospheric Chemistry, 79(3), 153–166. 10.1007/s10874-022-09434-3

Isidorov, V., & Jdanova, M. (2002). Volatile organic compounds from leaves litter. Chemosphere, 48(9), 975–979. 10.1016/s0045-6535(02)00074-7

Junginger, M., Seidl, R., & Müller, J. (2025). *BETA-FOR_SPZ_Deadwood_Inventory_2014/2024 (1.0)* [Dataset]. Zenodo. 10.5281/zenodo.15007373

Jüttner, F. (1986). Analysis of organic compounds (VOC) in the forest air of the Southern Black Forest. Chemosphere, 15(8), 985–992. 10.1016/0045-6535(86)90551-5

Kacic, P., Gessner, U., Hakkenberg, C. R., Holzwarth, S., Müller, J., Pierick, K., Seidel, D., Thonfeld, F., Torresani, M., & Kuenzer, C. (2025). Characterizing local forest structural complexity based on multi-platform and -sensor derived indicators. Ecological Indicators, 170, 113085. 10.1016/j.ecolind.2025.113085

Kern, C., Burton, J., Raymond, P., D’Amato, A., Keeton, W., Royo, A., Walters, M., Webster, C., & Willis, J. (2016). Challenges facing gap-based silviculture and possible solutions for mesic northern forests in North America. Forestry, 90. 10.1093/forestry/cpw024

Kern, C. C., Montgomery, R. A., Reich, P. B., & Strong, T. F. (2013). Canopy gap size influences niche partitioning of the ground-layer plant community in a northern temperate forest. Journal of Plant Ecology, 6(1), 101–112. 10.1093/jpe/rts016

Kern, C. C., Montgomery, R. A., Reich, P. B., & Strong, T. F. (2014). Harvest-Created Canopy Gaps Increase Species and Functional Trait Diversity of the Forest Ground-Layer Community. Forest Science, 60(2), 335–344. 10.5849/forsci.13-015

Kessler, A., & Kalske, A. (2018a). Plant Secondary Metabolite Diversity and Species Interactions. Annual Review of Ecology, Evolution, and Systematics, 49(1), 115–138. 10.1146/annurev-ecolsys-110617-062406

Kessler, A., & Kalske, A. (2018b). Plant Secondary Metabolite Diversity and Species Interactions. Annual Review of Ecology, Evolution, and Systematics, 49(1), 115–138. 10.1146/annurev-ecolsys-110617-062406

Krumm, F., Kraus, D., & Deutschland (Eds.). (2013). *Integrative approaches as an opportunity for the conservation of forest biodiversity*. European Forest Institute.

Lachat, T., Brang, P., Bolliger, M., Bollmann, K., Brändli, U.-B., Bütler, R., Herrmann, S., Schneider, O., & Wermelinger, B. (2019). Entstehung, Bedeutung und Förderung. *Merkbl. Prax*.

Lappalainen, H. K., Sevanto, S., Bäck, J., Ruuskanen, T. M., Kolari, P., Taipale, R., Rinne, J., Kulmala, M., & Hari, P. (2009). Day-time concentrations of biogenic volatile organic compounds in a boreal forest canopy and their relation to environmental and biological factors. Atmospheric Chemistry and Physics, 9(15), 5447–5459. 10.5194/acp-9-5447-2009

Leather, S. R., Baumgart, E. A., Evans, H. F., & Quicke, D. L. J. (2014). Seeing the trees for the wood – beech ( *Fagus sylvatica* ) decay fungal volatiles influence the structure of saproxylic beetle communities. Insect Conservation and Diversity, 7(4), 314–326. 10.1111/icad.12055

Lira Dyson, B., & Bässler, C. (2025). *BETA-FOR_SP7_Deadwood_fruitbodies_2023_2024 (1.0)* [Dataset]. Zenodo. 10.5281/zenodo.16368089

MacArthur, R. H., & MacArthur, J. W. (1961). On Bird Species Diversity. Ecology, 42(3), 594–598. 10.2307/1932254

Mäki, M., Aaltonen, H., Heinonsalo, J., Hellén, H., Pumpanen, J., & Bäck, J. (2019). Boreal forest soil is a significant and diverse source of volatile organic compounds. Plant and Soil, 441. 10.1007/s11104-019-04092-z

Mäki, M., Mali, T., Hellén, H., Heinonsalo, J., Lundell, T., & Bäck, J. (2021). Deadwood substrate and species-species interactions determine the release of volatile organic compounds by wood-decaying fungi. Fungal Ecology, 54, 101106. 10.1016/j.funeco.2021.101106

Massó Estaje, C., Rothacher, J., Vujić, A., Miličić, M., Chao, A., Mitesser, O., Claßen, A., & Steffan-Dewenter, I. (2025). *Experimental enhancement of structural heterogeneity in forest landscapes promotes multidimensional hoverfly diversity*.

Monard, C., Caudal, J.-P., Cluzeau, D., Le Garrec, J.-L., Hellequin, E., Hoeffner, K., Humbert, G., Jung, V., Le Lann, C., & Nicolai, A. (2021). Short-Term Temporal Dynamics of VOC Emissions by Soil Systems in Different Biotopes. Frontiers in Environmental Science, 9, 650701. 10.3389/fenvs.2021.650701

Müller, J. (2025). *BETA-FOR_SP9_Coleoptera_2022/2023 (1.0)* [Dataset]. Zenodo. 10.5281/zenodo.15396245

Müller, J., & Brandl, R. (2009). Assessing biodiversity by remote sensing in mountainous terrain: The potential of LiDAR to predict forest beetle assemblages. Journal of Applied Ecology, 46(4), 897–905. 10.1111/j.1365-2664.2009.01677.x

Müller, J., Scherer-Lorenzen, M., Ammer, C., Eisenhauer, N., Seidel, D., Schuldt, B., Biedermann, P., Schmitt, T., Künzer, C., Wegmann, M., Cesarz, S., Peters, M., Feldhaar, H., Steffan-Dewenter, I., Claßen, A., Bässler, C., von Oheimb, G., Fichtner, A., Thorn, S., & Weisser, W. (2022). BETA-FOR: Enhancing the structural diversity between patches for improving multidiversity and multifunctionality in production forests. Proposal for DFG Research Unit FOR 5375BETA-FOR: Erhöhung der strukturellen Diversität zwischen Waldbeständen zur Erhöhung der Multidiversität und Multifunktionalität in Produktionswäldern. Antragstext für die DFG Forschungsgruppe FOR 5375: β\(_4\) : Proposal for the 1st phase (2022-2026) of the DFG Research Unit FOR 5375/1 (DFG Forschergruppe FOR 5375/1 – BETA-FOR), Fabrikschleichach, October 2021 (p. 11531 KB, 210 pages) [Application/pdf]. Universität Würzburg. 10.25972/OPUS-29084

Noe, S. M., Hüve, K., Niinemets, Ü., & Copolovici, L. (2012). Seasonal variation in vertical volatile compounds air concentrations within a remote hemiboreal mixed forest. Atmospheric Chemistry and Physics, 12(9), 3909–3926. 10.5194/acp-12-3909-2012

Otieno, M., Karpati, Z., Peters, M. K., Duque, L., Schmitt, T., & Steffan-Dewenter, I. (2023). Elevated ozone and carbon dioxide affects the composition of volatile organic compounds emitted by Vicia faba (L.) and visitation by European orchard bee (Osmia cornuta). PLOS ONE, 18(4), e0283480. 10.1371/journal.pone.0283480

Ottensmann, M., Stoffel, M., Nichols, H., & Hoffman, J. (2018). GCalignR: An R package for aligning gas-chromatography data for ecological and evolutionary studies. PLOS ONE, 13, e0198311. 10.1371/journal.pone.0198311

Peñuelas, J. & Llusià, J. (2003). BVOCs: Plant defense against climate warming? Trends in Plant Science, 8(3), 105–109. 10.1016/S1360-1385(03)00008-6

Petersen, R., Holst, T., Mölder, M., Kljun, N., & Rinne, J. (2023). Vertical distribution of sources and sinks of volatile organic compounds within a boreal forest canopy. Atmospheric Chemistry and Physics, 23(13), 7839–7858. 10.5194/acp-23-7839-2023

Pierick, K., & Ammer, C. (2025). *BETA-FOR_SPZ_Tree inventory (1.0)* [Dataset]. Zenodo. Pierick, K., & Ammer, C. (2025). BETA-FOR_SPZ_Tree inventory (1.0) [Data set]. Zenodo. 10.5281/zenodo.15102089

Pollmann, J., Ortega, J., & Helmig, D. (2005). Analysis of Atmospheric Sesquiterpenes: Sampling Losses and Mitigation of Ozone Interferences. Environmental Science & Technology, 39(24), 9620–9629. 10.1021/es050440w

R Core Team. (2025). *R: A Language and Environment for Statistical Computing* [R]. R Foundation for Statistical Computing. https://www.R-project.org/

Raguso, R. A. (2008). Wake Up and Smell the Roses: The Ecology and Evolution of Floral Scent. Annual Review of Ecology, Evolution, and Systematics, 39(Volume 39, 2008), 549–569. 10.1146/annurev.ecolsys.38.091206.095601

Ramirez, K. S., Lauber, C. L., & Fierer, N. (2010). Microbial consumption and production of volatile organic compounds at the soil-litter interface. Biogeochemistry, 99(1/3), 97– 107.

Randlkofer, B., Obermaier, E., Hilker, M., & Meiners, T. (2010). Vegetation complexity— The influence of plant species diversity and plant structures on plant chemical complexity and arthropods. Basic and Applied Ecology, 11(5), 383–395. 10.1016/j.baae.2010.03.003

Riches, M. (2023, May 3). The Untold Story of the Understory: Direct Emission of Volatile Organic Compounds from Understory Species of a Ponderosa Pine Forest and Their Potential Role in Atmospheric Chemistry. 35th Conference on Agricultural and Forest Meteorology/14th Fire and Forest Meteorology Symposium/Sixth Conference on Atmospheric Biogeosciences. https://ams.confex.com/ams/35AF14F6BG/webprogram/Paper423943.html

Ringsdorf, A., Edtbauer, A., Holanda, B., Poehlker, C., Sá, M. O., Araújo, A., Kesselmeier, J., Lelieveld, J., & Williams, J. (2024). Investigating carbonyl compounds above the Amazon rainforest using a proton-transfer-reaction time-of-flight mass spectrometer (PTR-ToF-MS) with NO^+^ chemical ionization. Atmospheric Chemistry and Physics, 24(20), 11883–11910. 10.5194/acp-24-11883-2024

Rothacher, J., Hagge, J., Bässler, C., Brandl, R., Gruppe, A., & Müller, J. (2023). Logging operations creating snags, logs, and stumps under open and closed canopies promote stand-scale beetle diversity. Forest Ecology and Management, 540, 121022. 10.1016/j.foreco.2023.121022

Schieweck, A. (2018). Analytical procedure for the determination of very volatile organic compounds (C3–C6) in indoor air.

Schuman, M. C. (2023). Where, When, and Why Do Plant Volatiles Mediate Ecological Signaling? The Answer Is Blowing in the Wind. Annual Review of Plant Biology, 74(1), 609–633. 10.1146/annurev-arplant-040121-114908

Schuman, M. C., Valim, H. A., & Joo, Y. (2016). Temporal Dynamics of Plant Volatiles: Mechanistic Bases and Functional Consequences. In J. D. Blande & R. Glinwood (Eds.), Deciphering Chemical Language of Plant Communication (pp. 3–34). Springer International Publishing. 10.1007/978-3-319-33498-1_1

Schwarz, R. (2025). *Inconsistent short-term effects of enhanced structural complexity on soil microbial properties across German forests*.

Steiger, S., Schmitt, T., & Schaefer, H. M. (2011). The origin and dynamic evolution of chemical information transfer. Proceedings of the Royal Society B: Biological Sciences, 278(1708), 970–979. 10.1098/rspb.2010.2285

Stein, A., & Kreft, H. (2015). Terminology and quantification of environmental heterogeneity in species-richness research. Biological Reviews, 90, 815. 10.1111/brv.12135

Sulzer, M., Brzozon, J., Christen, A., Dedden, L., Dormann, C., Dumberger, S., Frey, Y., Gassilloud, M., Göritz, A., Grote, R., Haberstroh, S., Kattenborn, T., Kremer, L., Kreuzwieser, J., Kühnhammer, K., Lang, F., Lee, H., Müller, J., Schack-Kirchner, H., … Werner, C. (2025). The ECOSENSE forest – enriching tower-based flux measurements of carbon and water exchange with novel distributed sensor networks. ARPHA Conference Abstracts, 8. 10.3897/aca.8.e149267

Tong, R., Ji, B., Wang, G. G., Lou, C., Ma, C., Zhu, N., Yuan, W., & Wu, T. (2024). Canopy gap impacts on soil organic carbon and nutrient dynamic: A meta-analysis. Annals of Forest Science, 81(1), 12. 10.1186/s13595-024-01224-z

Uhl, B., Krah, F.-S., Baldrian, P., Brandl, R., Hagge, J., Müller, J., Thorn, S., Vojtech, T., & Bässler, C. (2022). Snags, logs, stumps, and microclimate as tools optimizing deadwood enrichment for forest biodiversity. Biological Conservation, 270, 109569. 10.1016/j.biocon.2022.109569

Wickham, H., Averick, M., Bryan, J., Chang, W., McGowan, L. D., François, R., Grolemund, G., Hayes, A., Henry, L., Hester, J., Kuhn, M., Pedersen, T. L., Miller, E., Bache, S. M., Müller, K., Ooms, J., Robinson, D., Seidel, D. P., Spinu, V., … Yutani, H. (2019). Welcome to the Tidyverse. Journal of Open Source Software, 4(43), 1686. 10.21105/joss.01686

Yáñez-Serrano, A. M., Nölscher, A. C., Bourtsoukidis, E., Gomes Alves, E., Ganzeveld, L., Bonn, B., Wolff, S., Sa, M., Yamasoe, M., Williams, J., Andreae, M. O., & Kesselmeier, J. (2018). Monoterpene chemical speciation in a tropical rainforest:variation with season, height, and time of dayat the Amazon Tall Tower Observatory (ATTO). Atmospheric Chemistry and Physics, 18(5), 3403–3418. 10.5194/acp-18-3403-2018

